# CpG methylation and methionine metabolism account for phenotypic bifurcation of HIV transcription: a unique case of a pure epigenetic phenomenon

**DOI:** 10.64898/2026.08.10.743899

**Authors:** Nisha Gautam, Ankita Rai, Jakub Dopierała, Aleksandra Bryner, Marina Lusic, Heng-Chang Chen (陳珩昌)

**Author notes:** Equal contribution (Nisha Gautam, Heng-Chang Chen (陳珩昌)).

## Abstract

HIV transcription is characterized by its stochastic nature, which plays a pivotal role in determining the fate of a provirus—active replication or latent infection—and is therefore a critical determinant of the HIV latency establishment. Building upon our previous work, we identified a unique phenotype of stochastic HIV transcription in a Jurkat T cell clonal model harboring a single lentiviral-based vector, herein referred to as the HIV transcription-sensitized model. A defining feature of this cellular model is that the turnover of HIV transcription shows elevated frequency—a phenomenon designated phenotypic bifurcation—suggesting that, under certain conditions, the regulation of stochastic HIV transcription can be a pure epigenetic phenomenon. In continuation of and to further substantiate this premise, the present study characterizes the contributions of epigenetic regulation of CpG methylation, methionine metabolism that coordinates cell cycle events, and HIV antisense transcription to this phenomenon. This work adds direct causal evidence to the hypothesis that a potential lag prior to the entry of the G2 phase in the bifurcated state of low HIV transcription may serve as one of the underlying mechanisms that lead to the high CpG methylation level compared with that measured in the state of high HIV transcription, contributing to the cyclical turnover of phenotypic bifurcation of HIV transcription.

## Introduction

Sporadic and episodic manifestations of atypical phenotypes of HIV transcription—including phenotypic bifurcation^1,2^, deep latency^3,4^, and residual HIV transcription in the presence of latency-promoting agents (LPAs)^5^—have been documented; however, the underlying mechanisms governing these phenomena remain poorly understood. HIV phenotypic bifurcation was first reported by the study from Weinberger and colleagues^1^ in the context of an *in vitro* HIV bulk infection model, wherein a distinct subset of infected cells harboring lentiviral-based proviral vectors exhibited a bimodal distribution of GFP expression, characterized by the simultaneous presence of cells with markedly elevated and near-undetectable fluorescence levels. Importantly, this phenotypic bifurcation was found to correlate with specific integration site (IS) characteristics: proviruses exhibiting bifurcated expression were preferentially integrated in genomic loci proximal to human endogenous retroviral long terminal repeats, relative to those lacking this phenotype^1^. Subsequent investigations have further corroborated the influence of position effect variegation on genome-wide fluctuations in HIV transcriptional activity^5,6^, collectively underscoring the synergistic interplay between epigenetic modifications and proviral IS in governing stochastic HIV transcription.

Unlike transcriptional heterogeneity arising from proviral integration across distinct chromosomal loci, we previously demonstrated that phenotypic bifurcation of HIV transcription can emerge from a single, defined proviral IS, as established using clonal cell populations^2^. This cellular clone is named the HIV transcription sensitized model in this manuscript. This cellular system harbors a proviral IS previously mapped to chromosomal position 110,913,078 base pairs (bp) on chromosome 13 (chr13), residing within the protein-coding gene ankyrin repeat domain 10 (ANKRD10), in proximity to short interspersed nuclear element (SINE) and Alu repetitive elements^2^. A defining feature of this model is the reproducible, temporally constrained transition in the proportion of GFP-expressing cells—from a state characterized by an abundance of cells exhibiting high-intensity GFP fluorescence (hereafter referred to as GFP_Bright_) to one dominated by cells displaying markedly diminished fluorescence (hereafter referred to as GFP_Dim_)—a process that typically unfolds over approximately one week^2^. Given that this bifurcated transcriptional behavior originates from an identical proviral context, we postulate that the mechanisms underlying this phenomenon are predominantly epigenetic in nature.

## Results

### A higher level of host genomic CpG methylation was detected in GFP_Dim_

To quantify the CpG methylation level between GFP_Bright_ and GFP_Dim_, we FACS-sorted the HIV transcription sensitized model based on GFP expression (GFP_Bright_ versus GFP_Dim_), followed by immediate isolation of genomic DNA subjected to reduced representation bisulfite sequencing (RRBS)^7^, respectively. Two biological replicates were conducted for each GFP_Bright_ and GFP_Dim_. Although a discrepancy was observed between duplicates, a clear separation between GFP_Bright_ and GFP_Dim_ was observed (**Figure 1A**) and confirmed by the correlation coefficient matrix based on %methylation scores computed by methylKit^8^ (**Figure S1A**) and the methylation level (i.e., beta value, see **STAR Methods**) between replicates (**Figure S1B** and **S1C**). We further computed differential methylation regions (DMRs) from GFP_Bright_ versus Jurkat T cells (**Figure 1B**) and GFP_Dim_ versus Jurkat T cells (**Figure 1C**), followed by annotation to the human genome (hereinafter, differential methylation genes, DMGs) (see **STAR Methods**). 692 high and 1,687 low DMGs were retrieved in GFP_Bright_ (**Figure 1B**); 638 high and 1,708 low DMGs in GFP_Dim_ (**Figure 1C**) against the RRBS output from infection-free Jurkat T cells. 105 high DMGs are unique to GFP_Bright_, and 51 are unique to GFP_Dim_ (**Figure 1D**). 132 low DMGs are unique to GFP_Bright_, and 111 are unique to GFP_Dim_ (**Figure 1E**). While directly computing DMGs between GFP_Bright_ and GFP_Dim_, 386 DMGs were retrieved from GFP_Dim_, and 641 DMGs were retrieved from GFP_Bright_ (**Figure 1F**). Altogether, these findings exemplify that different methylation profiles can be introduced by bifurcated HIV transcription resulting from an identical provirus.

**Figure 1.**
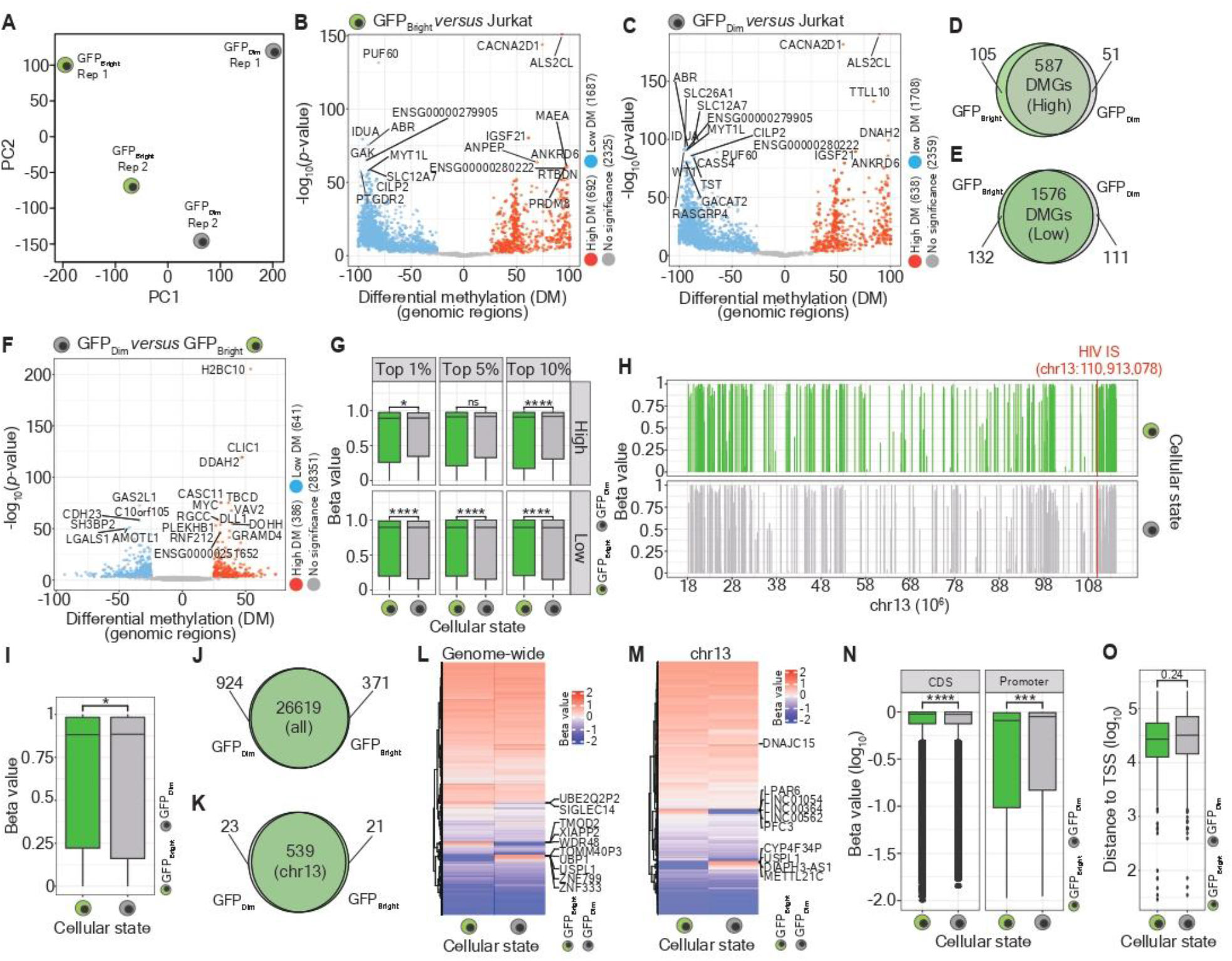
Discrete CpG methylation profiles between GFP_Bright_ and GFP_Dim_. (**A**) Principal component analysis (PCA) plot representing the discrepancy of the CpG methylation coverage between GFP_Bright_ and GFP_Dim_ offspring populations. (**B**, **C**) Volcano plots representing differential methylation regions (DMRs) retrieved from GFP_Bright_ (**B**) and GFP_Dim_ (**C**) offspring populations. Differential methylation was computed against methylation signals detected from virus-free Jurkat T cells. Dots marked in red indicate high differential methylation (DM); dots marked in blue indicate low DM; dots marked in grey indicate no significance. Numbers of measured DMRs are labeled in parentheses. 30% of non-significant DMRs were plotted. (**D**, **E**) Venn diagrams representing unique and common differently methylated genes (DMGs), separating high DM (**D**) from low DM (**E**), between GFP_Bright_ and GFP_Dim_. (**F**) Volcano plot representing DMRs between GFP_Bright_ and GFP_Dim_. Differential methylation was computed using DM computed from GFP_Dim_ against that computed from GFP_Bright_. Dots marked in red indicate high differential methylation (DM); dots marked in blue indicate low DM; dots marked in grey indicate no significance. Numbers of measured DMRs are labeled in parentheses. 30% of non-significant DMRs were plotted. (**G**) Box plot representing the CpG methylation level (i.e., beta value) across the top 1%, 5%, and 10% high and low methylation loci between GFP_Bright_ and GFP_Dim_. Facets on the x-axis separate the ranking of ^m^CpG loci; facets on the y-axis separate the methylation level (high versus low). The box marked in green refers to GFP_Bright,_ and the box marked in grey refers to GFP_Dim_. Significance levels are denoted as follows: ns for no significance, \**p* 0.05, \*\*\*\**p* 0.0001. (**H**) Line plots representing ^m^CpG sites and their intensity throughout chr13. Single provirus IS is highlighted in red. (**I**) Box plot representing the quantitative methylation level at chr13 between GFP_Bright_ and GFP_Dim_. The box marked in green refers to GFP_Bright,_ and the box marked in grey refers to GFP_Dim_. Significance levels are denoted as follows: ns for no significance, \**p* 0.05. (**J**, **K**) Venn diagrams representing host genes harboring CpG methylation signal between GFP_Bright_ and GFP_Dim_ at the genome-wide level (**J**) or at chr13 (**K**). (**L**, **M**) Clustering heatmap representing host genes harboring CpG methylation signal between GFP_Bright_ and GFP_Dim_ at the genome-wide level (**L**) or at chr13 (**M**). The top 10 genes with diverse CpG methylation levels between the two subpopulations are labeled on the right-hand side of the heatmap. (**N**) Box plots representing the CpG methylation level annotated in coding sequences (CDS) or the promoter regions between GFP_Bright_ and GFP_Dim_ at chr13. The x-axis is on a logarithmic scale. The box marked in green refers to GFP_Bright,_ and the box marked in grey refers to GFP_Dim_. Significance levels are denoted as follows: ns for no significance, \*\*\**p* 0.01, \*\*\*\**p* 0.0001. (**O**) Box plot representing the distance of ^m^CpG loci towards the closest transcriptional start sites (TSS) between GFP_Bright_ and GFP_Dim_ at chr13. The box marked in green refers to GFP_Bright_ and the box marked in grey refers to GFP_Dim_. The x-axis is on a logarithmic scale. Significance levels are denoted as follows: ns for no significance, \*\*\**p* 0.01, \*\*\*\**p* 0.0001.

We further performed gene ontology (GO) and Kyoto Encyclopedia of Genes and Genomes (KEGG) pathway over-representation analysis using DMGs. While using DMGs retrieved from the comparison between GFP_Bright_ and GFP_Dim_, 83 GO terms in Biological Process (BP) were enriched in high DMGs (n = 441) (**Table S1**), and 47 terms were associated with low DMGs (n = 596) (**Table S2**); the top 20 enriched GO terms respective to high and low DMGs were plotted in **Figure S2A**. 13 KEGG pathways were over-represented in high DMGs (**Table S3**), whereas no over-represented pathways were associated with low DMGs. While using DMGs separately retrieved from GFP_Bright_ versus Jurkat cells and GFP_Dim_ versus Jurkat cells, 0 and 154 (**Table S4**) GO terms in Biological Process (BP) were enriched from high (n = 563) and low (n = 1,308) DMGs in GFP_Bright_, respectively; 0 and 254 (**Table S5**) GO terms in Biological Process (BP) were enriched from high (n = 517) and low (n = 1,328) DMGs in GFP_Dim_, respectively. The top 20 enriched GO terms respective to each scenario were plotted in **Figure S2B**. 16 (**Table S6**) and 19 (**Table S7**) KEGG pathways were over-represented using low DMGs in GFP_Bright_ and GFP_Dim_, respectively. No pathways were enriched in high DMGs from both sets of cells. This observation suggested that relative to DMGs in GFP_Bright_, a higher functional similarity is shared among DMGs in GFP_Dim_.

While quantifying the beta value between GFP_Bright_ and GFP_Dim_, we ranked the top 1%, 5%, and 10% loci with the highest or the lowest beta values (**Figure 1G**). Relative to GFP_Bright_, we observed a relatively high beta value in GFP_Dim_ across the top 1%, 5%, and 10% ranked loci with the highest beta values. In contrast, a reverse pattern was observed across the top 1%, 5%, and 10% ranked loci with the lowest beta values (**Figure 1G**). This observation suggests a relatively high methylation level in GFP_Dim_. Next, we plotted the methylation level landscape throughout chr13, in which the provirus was mapped (chr13:100,913,078), and observed more intense methylation signals surrounding HIV IS in GFP_Dim_ than those in GFP_Bright_ (**Figure 1H**). The overall methylation level throughout chr13 in GFP_Dim_ also showed a significant increase compared with that measured in GFP_Bright_ (**Figure 1I**), confirming the potential role of CpG methylation in regulating the HIV phenotypic bifurcation in this clone. No difference in differential methylation (DM) values between GFP_Bright_ and GFP_Dim_ at the genome-wide scale (**Figure S3A**) and at chr13 (**Figure S3B**) was observed. While ranking the top 1%, 5%, 10%, 20%, and 30% of high and low DMGs, an increase in DM values was observed in ranked low DMGs in GFP_Bright_ compared to those measured in GFP_Dim_ (**Figure S3C**); a minor or no difference in ranked high DMGs between GFP_Bright_ and GFP_Dim_ was shown (**Figure S3C**).

We annotated loci with RRBS-detected CpG methylation to respective genes: 569 genes are unique to GFP_Bright_ (**Figure 1J**), in which 21 unique genes are present at chr13 (**Figure 1K**), and 924 genes are unique to GFP_Dim_ (**Figure 1J**), in which 23 unique genes are present at chr13 (**Figure 1K**). While pairwise comparing beta values between GFP_Bright_ and GFP_Dim_ at the genome-wide scale (**Figure 1L**) or at chr13 (**Figure 1M**), a subset of the genes demonstrated diverse CpG methylation intensities. At the genome-wide scale, we observed zinc finger protein 333 (ZNF333), which has been previously reported to determine the trans CpG signature of single-nucleotide polymorphism rs6511961^9^.

The genes observed at chr13 were found to be associated with DNA methylation (CDC16, DNAJC15, METTL21C, and LPAR6), ubiquitination and RNA processing (USPL1), and long non-coding RNA and antisense transcripts (LINC01054, LINC00364, LINC00562, and DIAPH3-AS1). It is important to stress that DIAPH3-AS1 (DIAPH3 antisense RNA 1) has been reported to be involved in N^6^-methyladenosine (m^6^A) methylation^10^, and METTL21C (methyltransferase 21C, AARS1 lysine) has been reported to impose methylation at alanine tRNA synthetase^11^. Whether phenotypic bifurcation is mechanistically governed via methylation at the RNA level requires further investigation. Nevertheless, our findings signify the importance of CpG methylation in regulating stochastic HIV transcription and open an avenue of research on HIV phenotypic bifurcation, where epitranscriptomic modification (i.e., m^6^A modification) might also be involved. Lastly, we observed that a higher level of CpG methylation accumulated in coding sequence regions and in the vicinity of the promoter regions in GFP_Dim_ than in GFP_Bright_ (**Figure 1N**). Although a tendency that methylated CpG (^m^CpG) sites are more distal to transcriptional start sites in GFP_Dim_ than in GFP_Bright_ was observed (**Figure 1O**), no statistical significance was obtained (**Figure 1O**).

### A low HIV transcriptional state in GFP_Dim_ was associated with the abundance of HIV antisense transcripts

The consecutive competition between HIV sense and antisense transcription in this cellular model has been previously characterized^2^. In this work, we first performed reverse transcription (RT), followed by quantitative PCR (qPCR) targeting the regions of HIV 5’ long terminal repeat (5’LTR) and GFP, and observed a higher viral expression level in GFP_Bright_ and GFP_Dim_ (**Figure 2A**), confirming the transcriptional discrepancy between both subpopulations. To further examine whether HIV antisense transcripts (AST), which are known to be capable of competing its sense RNA transcription^2,12–14^ and most likely contributing to maintaining HIV latency^14^, also play a role in governing phenotypic bifurcation in our present cellular model, we performed strand-specific RT-qPCR with DNaseI-treated RNA isolated from GFP_Bright_ and GFP_Dim_, independently and observed a higher level of antisense transcription in GFP_Dim_ than that measured in GFP_Bright_ (**Figure 2B**); a revised pattern in sense transcripts (ST) was observed, although no statistical significance was obtained (**Figure 2B**). We further treated siRNAs against either ST—referred to as siRNA HCP#400—or AST— referred to as siRNAs HCP#397 and HCP#398—in GFP_Dim_ (**Figure 2C**) to validate the impact of AST in suppressing HIV transcription in GFP_Dim_. The siRNA HCP#399 was added to the experiment as a control, given its identical strand synthesis direction to ST and the absence of AST in the genomic region tagged by this siRNA (**Figure 2C**). A modest increase in the percentage of GFP-positive cells was measured in AST-depleted GFP_Dim_ compared with ST-depleted GFP_Dim_ and a control group (**Figure 2D**). A more obvious effect was observed when strand-specific RT-qPCR was conducted (**Figure 2E** and **2F**): a sharp increase in sense transcription was measured when AST were depleted (**Figure 2E**), whereas minimal and comparable sense transcription between ST-depleted GFP_Dim_ and a control group was observed (**Figure 2E**). Undetectable antisense transcription in AST-depleted GFP_Dim_ confirms the clean depletion of AST using designed siRNAs (**Figure 2F**). The absence of higher antisense transcription levels in GFP_Dim_ with depleted ST might result from the low baseline amount of AST, rendering any enrichment hard to detect (**Figure 2F**). Altogether, this finding suggests that HIV AST may also participate in the regulatory machinery of HIV phenotypic bifurcation.

**Figure 2.**
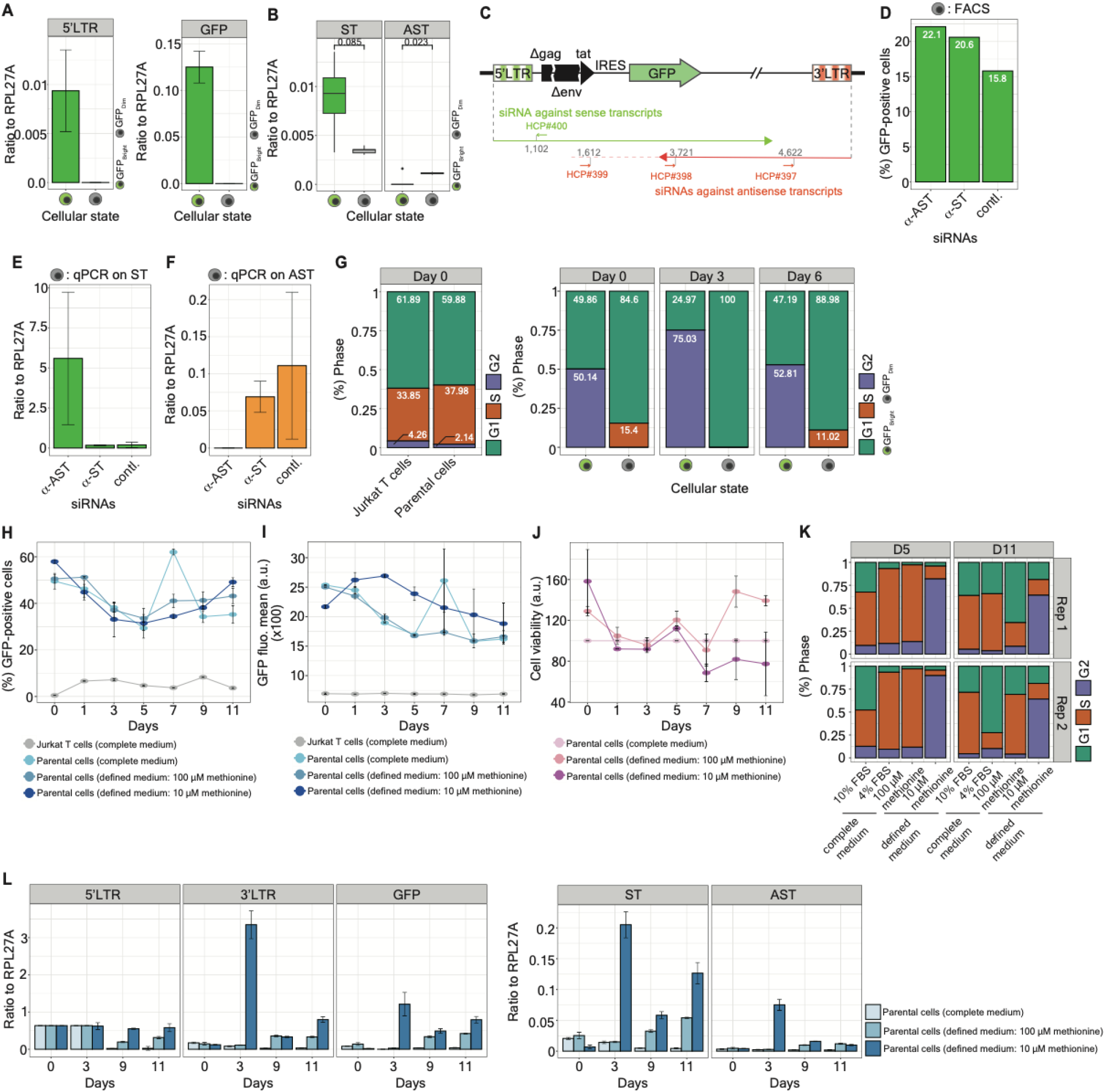
The characteristic features in antisense transcription and cell physiology of GFP_Bright_ and GFP_Dim_. (**A**) Bar chart representing the fold change in 5’LTR (left panel) and GFP (right panel) expression measured via RT-qPCR (ratio to RPL27A) between GFP_Bright_ and GFP_Dim_. The box marked in green refers to GFP_Bright,_ and the box marked in grey refers to GFP_Dim_. (**B**) Box plot representing the fold change in sense and antisense transcription measured via strand-specific RT-qPCR (ratio to RPL27A) between GFP_Bright_ and GFP_Dim_. Facets on the x-axis separate sense RNAs from antisense RNAs. The box marked in green refers to GFP_Bright,_ and the box marked in grey refers to GFP_Dim_. (**C**) Schematic representation of the design of siRNAs against sense transcripts—referred to as HCP#400—and antisense transcripts—referred to as HCP#397 and HCP#398. The siRNA HCP#399 serves as a control. (**D**) Bar chart representing the percentage of GFP-positive cells measured in ST-or AST-depleted GFP_Dim_, respectively, and in GFP_Dim_ treated with a control siRNA. (**E**, **F**) Bar charts representing the fold change in sense (**E**) and antisense transcription (**F**) measured via strand-specific RT-qPCR (ratio to RPL27A) in ST- or AST-depleted GFP_Dim_, respectively, and in GFP_Dim_ treated with a control siRNA. (**G**) Stacked bar graphs representing the proportion of the cells in the G1 (marked in green), S (marked in brown), or G2 (marked in purple) phase throughout the cell cycle. The Day 0 reading was taken on FACS-sorted GFP_Bright_ and GFP_Dim_ subjected to an immediate ice-cold 70% ethanol fixation. The measurement on Jurkat T cells and unsorted parental cells of GFP_Bright_ and GFP_Dim_ was performed in parallel. (**H**) Line plot representing the kinetics of the proportion of GFP-positive cells measured from unsorted parental cells grown in either complete medium or defined medium supplemented with different concentrations of methionine. Jurkat T cells grown in complete medium (grey line) serve as a control group. (**I**) Line plot representing the kinetics of the GPF fluorescence mean (arbitrary unit, a.u.) measured from unsorted parental cells grown in either complete medium or defined medium supplemented with different concentrations of methionine. Jurkat T cells grown in complete medium (grey line) serve as a control group. (**J**) Line plot representing the kinetics of cell viability (arbitrary unit, a.u.) measured from unsorted parental cells grown in either complete medium or defined medium supplemented with different concentrations of methionine. (**K**) Stacked bar graphs representing the proportion of the cells in the G1 (marked in green), S (marked in brown), or G2 (marked in purple) phase throughout the cell cycle. Unsorted parental cells were cultured in either the complete medium or the defined medium supplemented with 100 µM or 10 μM L-methionine. Measurements were performed in duplicate (Rep 1 and Rep 2) at d 5 and d 11 post-starvation. (**L**) Bar chart representing the fold change in 5’LTR, 3’LTR, GFP, ST, and AST (from the left-to right-hand side) expression measured via RT-qPCR or strand-specific RT-qPCR (ratio to RPL27A) at d 0, d 3, d 9, and d 11 post-starvation. DNaseI-treated RNA was isolated from unsorted parental cells grown in either complete medium or defined medium supplemented with different concentrations of methionine.

### The coordinate between cell cycle events and methionine metabolism regulates bifurcated states of HIV transcription

A question arising from the abovementioned observations regarding distinct CpG methylation levels between GFP_Bright_ and GFP_Dim_ is whether both offspring subpopulations also possess discrete cell cycle profiles and methionine metabolism. While delineating the cell cycle profile at three independent time points, the majority of the GFP_Dim_ remained in the G1 and S phases (**Figure 2G**), whereas partial cells in GFP_Bright_ shifted to the G2 phase, resulting in the absence of GFP_Bright_ in the S phase (**Figure 2G**). Profiles between GFP_Bright_ and GFP_Dim_ differ from those generated from infection-free Jurkat T cells and unsorted parental cells (**Figure 2G**). This finding led us to hypothesize that methionine metabolism, through its engagement with the cell cycle, regulates bifurcated HIV transcriptional states, whereby delayed G2 phase entry in GFP_Dim_ facilitates methyltransferase synthesis and maintenance—the fuel for methylation—during the S phase. To verify our hypothesis, we performed the longitudinal methionine restriction assay (see **STAR Methods**) and determined the kinetics of the proportion of GFP-positive cells using unsorted parental cells (**Figure 2H**). Under the methionine starvation condition (i.e., 10 µM L-methionine), the percentage of GFP-positive cells showed an obvious increase from d 5 to d 11 post-starvation (**Figure 2H**). Despite a reduction in GFP fluorescence mean across all settings, the drop was minimal under the methionine starvation condition (**Figure 2I**). A minor dose-dependent reduction in cell viability— about 77% of the cells remained alive—was observed at d 11 in the methionine starvation condition (**Figure 2J**). Upon repeating the cell cycle assay at d 5 and d 11 post-starvation, the majority of cells remained in the G2 phase (**Figure 2K**). In contrast, the majority of the cells cultured in the defined medium supplemented with 100 μM L-methionine and the complete medium were predominantly distributed in the S phase (**Figure 2K**). These findings suggest that methionine starvation may promote G2-phase entry, which in turn could contribute to the observed increase in the percentage of GFP-positive cells and GFP fluorescence mean (**Figure 2H** and **2I**). Lastly, we performed RT-qPCR and strand-specific RT-qPCR to confirm viral expression in the elevated proportion of GFP-positive cells under the methionine starvation condition (**Figure 2L**): 5’LTR, 3’ long terminal repeat (3’LTR), and GFP expression, as well as ST and AST abundances, showed an increase when methionine concentration is restricted (**Figure 2L**). Altogether, an important implication of these results is that the engagement of methionine metabolism with the cell cycle can constitute a conserved layer of the regulation of HIV phenotypic bifurcation.

### ^m^CpG sites proximal to HIV IS play a role in determining bifurcated states of HIV transcription

To rank the importance of the ^m^CpG loci (n = 66) and their targeting genes (n = 40) presented at chr13 in determining phenotypic bifurcation, we applied three machine learning-based models, including random forest (RF), XGBoost, and Pearson correlation, for the classification of bifurcated states in this model. Only methods based on RF and Pearson correlation enabled an appropriate classification (**Figure 3A**). We thus computed competence scores (see **STAR Methods**), reflecting the importance of ^m^CpG loci or ^m^CpG-targeted genes based on these two algorithms. At a locus level, the most important loci were chr13:113,830,863 (score: 0.897), followed by chr13:64,734,004 (score: 0.887), chr13:113,089,387 (score: 0.748), chr13:64,734,009 (score: 0.654), and chr13:20,998,701 (score: 0.542) (**Figure 3B** and **3C**). We observed a modest negative correlation between the distance of the ^m^CpG loci towards HIV IS and competence score (**Figure 3D**)—loci with a score over 0.5 (the top 5 ranked loci) were clearly separated from other loci (**Figure 3D**). At a gene level, the most important gene was ENSG00000283828 (score: 0.636), followed by MCF2L (score: 0.617), IRS2 (score: 0.509), and MTIF3 (score: 0.502) (**Figure 3E** and **3F**). These genes have been previously reported to be involved in the epigenetic regulation of CpG methylation in non-infectious diseases^15–17^. Superior classification performance introduced at a locus level suggests that resolution down to a nucleotide level may be required for the classification of phenotypic bifurcation. Moving forward, we stimulated 66 ^m^CpG sites identified at chr13 between GFP_Bright_ and GFP_Dim_ (see **STAR Methods**) and observed three loci that achieve statistical significance (**Figure 3G**). Among them, the ^m^CpG site at chr13:110,869,930 (#51) is proximal to the HIV IS when delineating all simulated levels of CpG methylation at chr13 (**Figure 3H**). The causal reasons for only a few ^m^CpG sites that reach statistical significance (**Figure 3G**) might be due to either epigenetic noise or an inadequate number of ^m^CpG sites in the simulation. Nevertheless, these findings suggest that the distance of ^m^CpG sites toward the HIV IS can dictate the mechanism driving phenotypic bifurcation.

**Figure 3.**
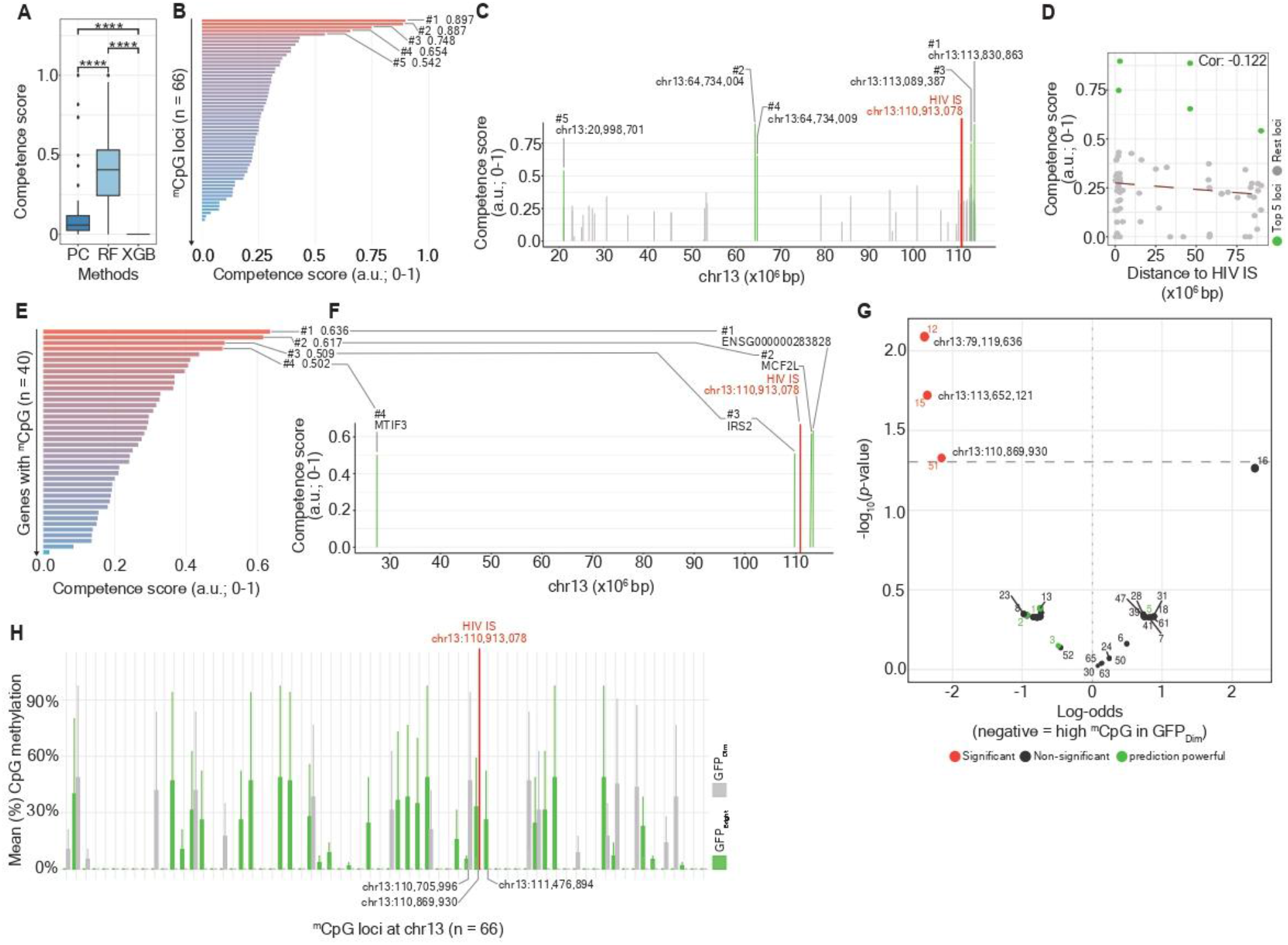
Ranking the importance of ^m^CpG loci and genes involved in HIV phenotypic bifurcation. (**A**) Box plot representing the competence of different machine learning-based models, including random forest (RF), XGBoost (XGB), and Pearson correlation (PC) for feature importance ranking. Significance levels are denoted as follows: ns for no significance, \*\*\*\**p* 0.0001. (**B**) Bar chart representing the ranking importance of 66 ^m^CpG loci (y-axis) at chr13. The x-axis indicates the competence score, ranging from 0 to 1 (1: most important; 0: no importance). (**C**) The landscape of 66 ^m^CpG loci at chr13. The top 5 most important loci responsible for phenotypic bifurcation are highlighted in green. Single provirus IS is highlighted in red. (**D**) Scatter plot representing the negative correlation between the distance of the ^m^CpG loci towards HIV IS and competence score. Dots marked in green indicate the top 5 most important ^m^CpG loci; the rest of the ^m^CpG loci were marked in grey. (**E**) Bar chart representing the ranking importance of 40 ^m^CpG genes (y-axis) at chr13. The x-axis indicates the competence score, ranging from 0 to 1 (1: most important; 0: no importance). (**F**) Schematic representation of the positions of the top 4 most important ^m^CpG genes responsible for phenotypic bifurcation is marked in green. Single provirus IS is highlighted in red. (**G**) Volcano plot representing the significance of individual CpG loci at chr13 based on the simulation of observed methylation levels between GFP_Bright_ and GFP_Dim_. The horizontal dashed line denotes the value of the negative decadic logarithm of a *p*-value equal to 0.05. The size of a dot denotes the value of the negative decadic logarithm of a *p*-value. Dots marked in red represent the loci with significant *p*-values; dots marked in black represent the loci with non-significant *p*-values; dots marked in green represent loci with competence scores greater than 0.5. (**H**) Bar chart representing simulated CpG methylation level across 66 loci throughout chr13 between GFP_Bright_ and GFP_Dim_. Bars marked in green refer to the CpG methylation level in GFP_Bright_; bars marked in green refer to the CpG methylation level in GFP_Dim_.

## Discussion

In this work, we showcase a correlation between host CpG methylation and bifurcated states of HIV transcription: higher methylation corresponds to the low transcription marked by GFP_Dim_ (**Figure 1**). This state is also associated with greater abundance of HIV AST (**Figure 2B**) and a delayed entry in the G2 phase relative to GFP_Bright_ (**Figure 2G**). Modeling and simulation results further show that ^m^CpG sites proximal to the provirus IS are more predictive of bifurcated states of HIV transcription than distal sites (**Figure 3**). Altogether, these results suggest that CpG methylation, methionine-linked cell cycle regulation, and antisense transcription can constitute a conserved layer to govern HIV phenotypic bifurcation.

HIV infections reprogram host immune cells epigenetically, including through changes in CpG methylation. Studies of immune cells from chronic HIV-infected individuals have revealed methylation signatures associated with antiretroviral therapy (ART)^18–21^, disease progression^22,23^, and advanced epigenetic aging^24,25^. Genome-wide CpG methylation changes, however, are detectable even during acute infections in individuals who initiate ART within three days after diagnosis^26^. The early initiation of ART can minimally restore acute HIV-related CpG methylation changes in monocytes at interferon-related genes, whereas no impact on the alteration of the CpG methylation profile in CD4+ T lymphocytes^26^. Zhang and colleagues similarly reported distinct CpG methylation patterns across five different immune cells—CD4+ T cells, CD8+ T cells, B cells, Natural Killer cells, and monocytes^27^—reinforcing that host CpG methylation and HIV pathogenesis are tightly linked from the early stage of infection.

A recent finding pointing out that HIV-related methylation changes more frequently affect low activity regions of the genome, including heterochromatin and quiescent regions^28^, resonates with earlier work showing that the latent reservoir in elite controllers are enriched in heterochromatin^4^—more than 90% of cytosine residues are hypermethylated in proximity to HIV IS preferentially situated in centromeric satellite DNA and in Krüppel-associated box domain-containing zinc finger genes, both of which are associated with heterochromatin features^4^. Three significant CpG sites—g04784635, cg13131185, and cg07189782—have also been repeatedly implicated in various HIV controllers, including elite controllers^29^. Our results extend this picture by implicating methionine metabolism and cell cycle events in CpG methylation differences that distinguish HIV phenotypic bifurcation.

While annotating the CpG methylation profile across the gene architecture, the observation that a higher CpG methylation level in the promoter region in GFP_Dim_ relative to GFP_Bright_ (**Figure 1N**) aligns with the current general principle that promoter-associated CpG methylation suppresses gene expression^30,31^. In contrast to the previous study stating TSS-proximal hypermethylation to latency, no significant difference in the distribution of ^m^CpG loci toward the closest TSS between GFP_Bright_ and GFP_Dim_ was observed using our clonal model (**Figure 1O**). This inconsistency can be further investigated when additional HIV transcription sensitized cellular models are available. Nevertheless, our result that ^m^CpG loci in the proximity to the HIV IS offer better prediction performance in terms of bifurcated states of HIV transcription (**Figure 3A–3D**) points to the discrete local chromatin context surrounding HIV IS as a likely contributor to phenotypic bifurcation. This finding resonates with the previous work on position effect variegation in determining HIV transcription^1,6^. Whether host CpG methylation alone is sufficient to drive phenotypic bifurcation, however, remains unresolved. Future investigations should pursue both the underlying mechanism behind the dynamic alteration of the CpG methylation level over time and a broader regulatory network in which host CpG methylation, viral proteins, and host cellular factors interact.

The discrete CpG methylation profiles between GFP_Bright_ and GFP_Dim_ raise the question of whether cellular metabolism underlies bifurcated states of HIV transcription. Cellular metabolism^32^, with particular emphasis on glycolysis^33,34^, redox and iron metabolism^34,35^, tryptophan metabolism^36^, and SAMHD1-mediated intracellular dNTP levels^37^, has been shown to shape HIV replication in CD4+ T cells. Hypermetabolic responses in acquired immunodeficiency syndrome (AIDS) patients have been described in the late 20^th^ century^38–40^, and methionine was identified early on as one of the rate-limiting amino acids for whole-body protein synthesis in AIDS patients^41^, underscoring its role in nutritional status. In yeasts and mammalian cells, limiting methionine triggers G1/S arrest—the S-adenosylmethionine (SAM) checkpoint—via disassembly of pre-replication complexes^42^ and cyclin-dependent kinase 2 hypophosphorylation^43,44^. Our result that GFP_Dim_ accumulates at the G1/S phases relative to GFP_Bright_ exemplifies a physiological difference between these two subpopulations (**Figure 2G**). We hypothesize that a delay before G2 entry in GFP_Dim_ enables the accumulation and maintenance of methyltransferase, subsequently producing the higher methylation level observed in GFP_Dim_. While the cell cycle dynamics of DNA methylation are not fully understood, our hypothesis aligns with the previous studies of DNA methyltransferase 1 (DNMT1) retention^45^ and increased methylation during the S phase^46^, and a lag in 5-methylcytosine (5mC) accumulation in early G2/M phases before peak levels of 5mC were observed^47^. Supporting this link, result from the methionine restriction assay conclusively evidences the profound involvement of methionine metabolism in the regulatory machinery of bifurcated states of HIV transcription (**Figure 2H** and **2I**)—methionine starvation, which triggers a reduction in the SAM-to-S-adenosylhomocysteine (SAH) ratio, subsequently halting the methylation capacity—can serve as one of the causal mechanisms behind the regulation of distinct CpG methylation levels between GFP_Bright_ and GFP_Dim_. Phenotypically, the observation that methionine starvation promotes G2-phase entry (**Figure 2K**) further strengthens our hypothesis. In mammalian cells, the serine/threonine kinase mTOR is responsible for the sensing of methionine and its derivates. Although methionine depletion has been reported to induce a SAMTOR (S-adenosylmethionine sensor upstream of mTORC1)-dependent inhibition of mTOR complex 1 (mTORC1)^48^, a well-validated regulator of lifespan and health in many organisms, the core sensors that sense methionine or SAM abundance in mammalian cells remain unclear. These questions will be addressed in further investigation of the coordination between SAM-checkpoint arrest at the G1 phase and HIV phenotypic bifurcation, or HIV transcription in general. It is noteworthy that the study reported by Yang and colleagues has characterized the contribution of methionine adenosyltransferase 2A (MAT2A)—the enzyme required to convert methionine to SAM under ATP consumption—to HIV latency via SAM-mediated one-carbon flux, as SAM is the primary methyl group donor and is required for epigenetic regulation and other methylation-controlled processes^49^.

Taken together, results presented in this work and previous findings underscore that the HIV life cycle is shaped by the metabolic state of the infected cell, reinforcing the importance of studying cellular metabolism alongside viral pathogenesis. Under the present conceptual framework of the HIV latency establishment, latency is more like a bypass product when CD4+ T cells enter quiescence. Although in other viruses, such as herpes viruses and human cytomegalovirus, their antisense gene product or antisense transcripts have been reported to be capable of promoting latency, whether HIV AST can also play an active role in programming HIV latency remains unknown. Our siRNA depletion experiments suggest that HIV AST regulate bifurcated states of HIV transcription (**Figure 2B**–**2F**). Consistency in elevated ST and AST abundances under the methionine starvation condition (**Figure 2L**) recaptures our previous finding of the simultaneous transcriptional pattern between ST-AST competition^2^, implying that consecutive ST-AST competition can be cooperative. Future experiments depleting AST in models of inducible versus deep latency will help clarify whether AST actively programmes proviral entry and relief from latency. Further studies are needed to elucidate the changes in methionine metabolic status throughout the cell cycle and to determine whether this amino acid contributes to the pathophysiology of HIV infection, epigenetic regulation, and the latency establishment in the context of bifurcated states of HIV transcription.

### Limitations of this study

One of the major limitations is a lack of methylation profiling throughout the 5’LTR, as HIV latency has been proposed to be linked to 5’LTR methylation. However, the mechanism by which CpG methylation on the HIV 5’LTR promotes HIV latent infections is presently debated. Some studies have reported high levels of CpG methylation in the promoter region of the HIV 5’LTR^50,51^; others have found this region to be unmethylated^52,53^ or to have low methylation level^54^. Methylation of the 3’LTR, which drives antisense transcription, is comparatively understudied. Therefore, a profound characterization of dynamic CpG methylation patterns on both HIV LTRs, accompanied by the cyclical turnover of bifurcated states of HIV transcription, and examination of their molecular interplay with critical factors involved in HIV transcription machinery, such as RNA polymerase II pausing and elongation, will provide more mechanistic insights into the dynamic regulation of fluctuations in HIV transcription and the latency establishment. Furthermore, it remains unclear how bifurcated transcriptional states of HIV respond to latency-reversing and latency-promoting agents, and whether proviral reactivation exhibits comparable phenotypic bifurcation governed by shared mechanisms and factors. Finally, despite distinct GO terms (**Figure S2**) and KEGG pathways (**Tables S3**, **S6**, and **S7**) enriched between GFP_Bright_ and GFP_Dim_, future work should also determine whether additional transcription factors or signaling pathways triggered by methionine starvation co-regulate HIV phenotypic bifurcation via the engagement of methionine metabolism with the cell cycle.

## Acknowledgements

We would like to thank Dr. hab. Marek Wagner (Innate Immunity Research Group, Łukasiewicz Research Network – PORT Polish Center for Technology Development) and Mr. Mateusz Marciniak (Innate Immunity Research Group, Łukasiewicz Research Network – PORT Polish Center for Technology Development) for sharing the cell proliferation assay protocol and lending Multi-Mode Microplate Reader (Agilent Technologies) for conducting the experiment. H.-C.C. acknowledges funding from the Narodowe Centrum Nauki (Sonata Bis Grant UMO-2022/46/E/NZ6/00022 and UMO-2024/06/Y/NZ6/00041).

## Author contributions

Conceptualization, H.-C.C.; methodology, H.-C.C., M.L., and N.G., A.R.; software, H.-C.C., formal analysis, H.-C.C., investigation, H.-C.C., N.G., A.R., J.D., and A.B.; resources, H.-C.C.; data curation, H.-C.C.; writing of original draft manuscript, H.-C.C., N.G., A.R.; writing, manuscript review and editing, H.-C.C., M.L., N.G., A.R.; visualization, H.-C.C.; supervision, H.-C.C., project administration, H.-C.C.; funding acquisition, H.-C.C.

## STAR Methods

## KEY RESOURCES TABLE

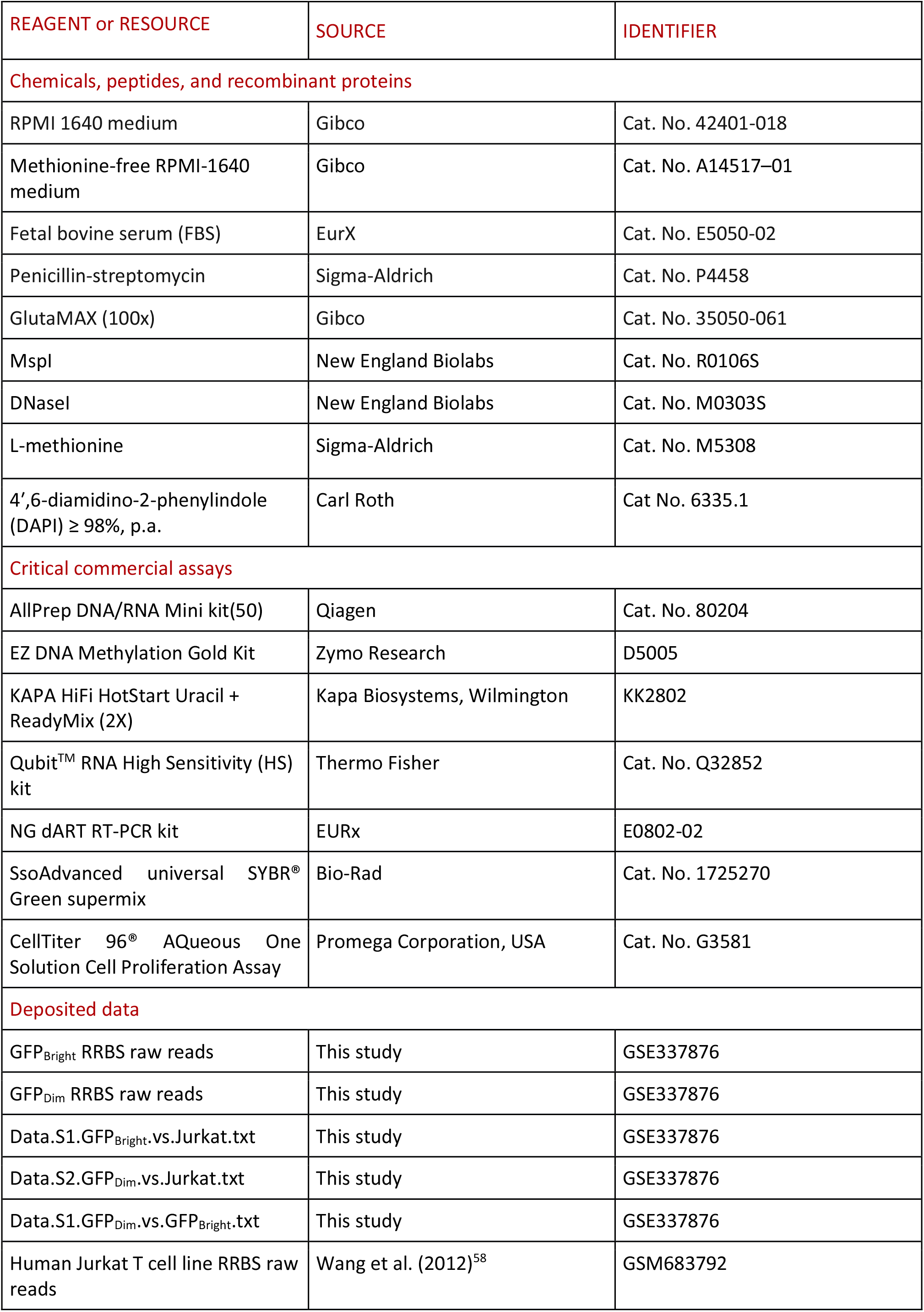
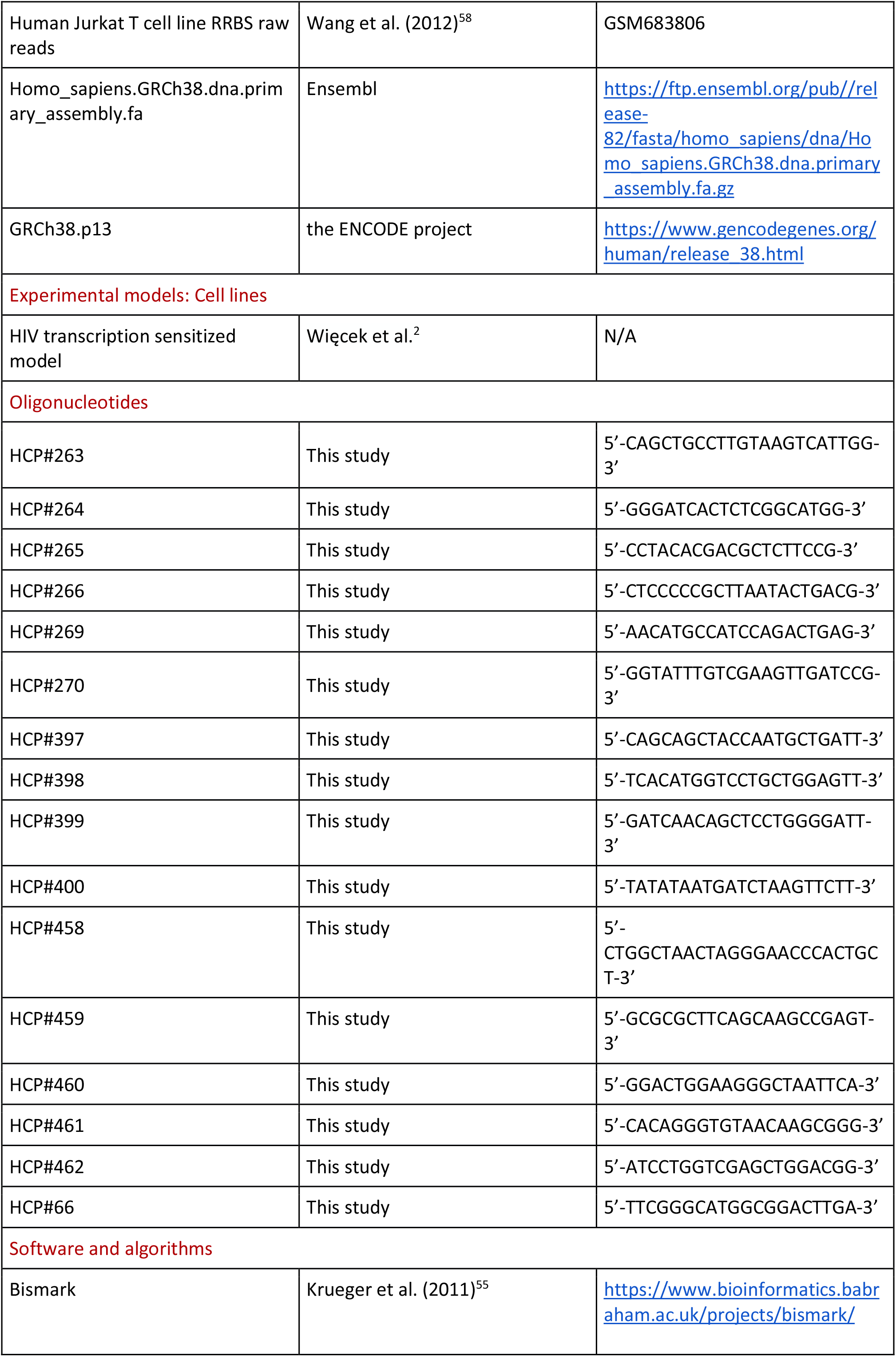
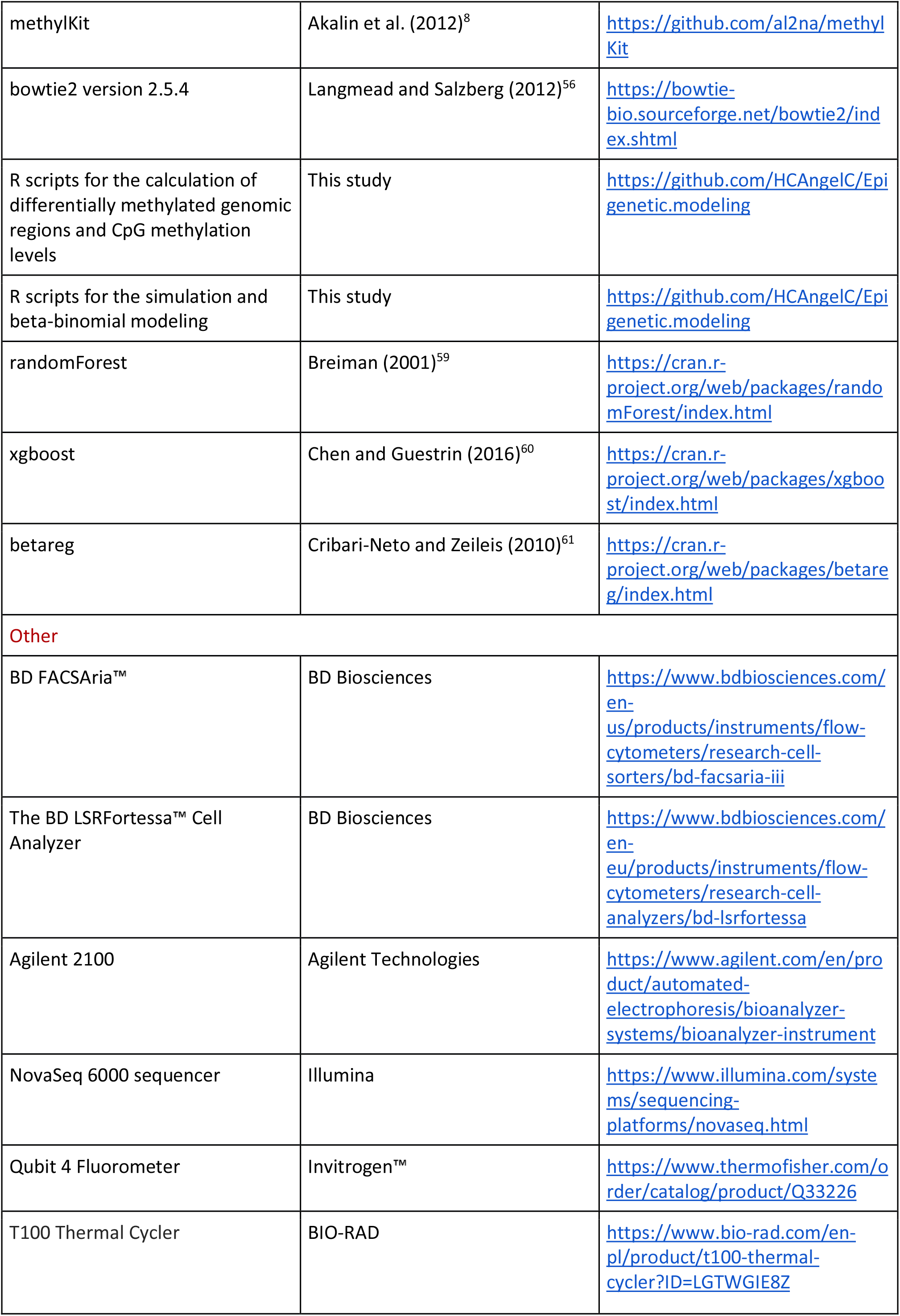
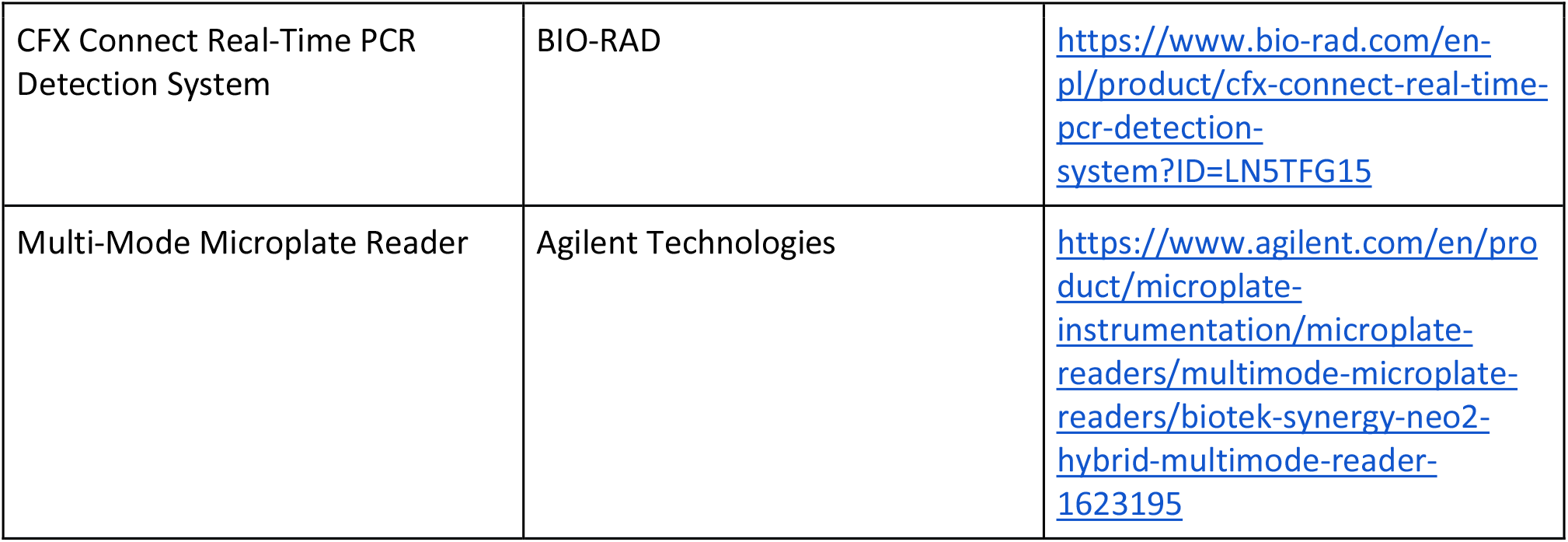

## RESOURCE AVAILABILITY

### Lead contact

Further information and requests for resources and reagents should be directed to and will be fulfilled by the lead contact, Heng-Chang Chen.

### Materials availability

This study did not generate new unique reagents.

### Data and code availability

#### Data availability

● The raw sequencing data generated during this study are available from Gene Expression Omnibus (GSE337876).
● Output files from the differential methylation analysis are available from Gene Expression Omnibus (GSE337876).
● Other analyzed data are provided in **Supplementary Tables**.

#### Code availability

● All code and scripts provided in this work are available on GitHub (https://github.com/HCAngelC/CpG.methylation.on.HIV.phenotypic.bifurcation).

Any additional information required to reanalyze the data reported in this paper is available from the lead contact upon request. This paper reports the original code.

## METHOD DETAILS

### Cell culture

The HIV transcription sensitized model^1^ was grown at 37 ◦C under a 95 % air and 5 % CO2 atmosphere, in RPMI 1640 medium (Gibco, Cat. No. 42401–018) supplemented with 10 % fetal bovine serum (FBS; EurX, Cat. No. E5050-02), 1 % penicillin-streptomycin (Sigma-Aldrich, Cat. No. P4458), and 1 % GlutaMAX (100x) (Gibco, Cat. No. 35050–061). Clonal cells were passaged every 2 d with a 1:5 dilution. To perform the methionine restriction assay, cells were cultured under three media conditions: (i) RPMI-1640 medium supplemented with 4% FBS, (ii) methionine-free RPMI-1640 medium (Gibco, Cat. No. A14517–01) supplemented with 4% FBS and 100 μM L-methionine (Sigma-Aldrich, Cat. No. M5308), and (iii) methionine-free RPMI-1640 medium supplemented with 4% FBS and 10 μM L-methionine.

### Preparation of RRBS sequencing libraries

GFP_Bright_ and GFP_Dim_ genomic DNA were isolated using the AllPrep DNA/RNA Mini Kit (50) (Qiagen, Cat. No. 80204) following the manufacturer’s instructions. 600 ng genomic DNA was digested with the methylation-insensitive restriction enzyme MspI (New England Biolabs, Cat. No. R0106S), followed by end-repair, A-tailing, and ligation with methylated adaptors. The DNA bisulfite conversion was performed using the EZ DNA Methylation Gold Kit (D5005, Zymo Research, Irvine, USA). DNA fragments were size-selected and amplified using the KAPA HiFi HotStart Uracil + ReadyMix (2X) (KK2802, Kapa Biosystems, Wilmington, USA). The library was analyzed for size distribution by Agilent 2100, and concentration was quantified using qPCR. The sequencing library was sequenced on an X plus flow cell with the PE150 strategy.

### Data analysis and bioinformatics

#### Analysis of RRBS readouts

Overall, sequencing reads from RRBS were mapped to Ensembl Homo_sapiens.GRCh38.dna.primary_assembly.fa, as the reference genome, using Bismark^55^ with the option ‘--genome’. The reference genome was indexed using bowtie2^56,57^ with the options ‘--path_to_aligner’ and ‘--verbose’ provided by Bismark. Two sets of RRBS sequencing reads conducted in Jurkat T cells (GSM683792 and GSM683806)^58^ were downloaded from NCBI Gene Expression Omnibus (GEO) and subjected to the Bismark analysis as a control of the infection-free condition. Mapped reads within an output file were selected and sorted by chromosome and the mapping position with the command line below.

~~~
$ grep -v \’^[[:space:]]*\@\’ mapped.sam | sort -k3,3 -k4,4n > mapped.sorted.sam
~~~

#### Calculation of differentially methylated genomic regions

The sorted output file can be executed using the R package methylKit^8^, followed by the computation of differentially methylated genomic regions (DMRs). Six different file lists, including (#1) GFP_Bright_ and GFP_Dim_, (#2) GFP_Bright_-Rep1, GFP_Bright_-Rep2, GFP_Dim_-Rep1, and GFP_Dim_-Rep2, (#3) GFP_Bright_ only, (#4) GFP_Bright_ and Jurkat T cells, (#5) GFP_Dim_ only, and (#6) GFP_Dim_ and Jurkat T cells, were prepared and called methylation percentage per base with the function processBismarkAln() and the argument ‘assembly = “hg38”’ provided in methylKit^8^. Samples in each file list were filtered based on read coverage with the function filterByCoverage() and the arguments ‘lo.count=10’, ‘lo.perc=NULL’, ‘hi.count=NULL’, and ‘hi.perc=99.9’. Samples within the same file list were merged with the function methylKit::unite(). Merged files from the file list #2 was used for generating the PCA plot with the function PCASamples(); merged files (#1, #4, and #6) were applied for the calculation of differentially methylated genomic regions (DMRs) with the function tileMethylCounts() and arguments ‘win.size=1000’ and ‘step.size=1000’, followed by the function calculateDiffMeth() with the argument ‘mc.cores = 2’. Output files of DMRs (#1, #4, and #6) can be found in Data S1–S3 (GSE337876) and were used for generating volcano plots.

DMRs were annotated to the human genome (GRCh38.p13, the ENCODE project) to obtain differentially methylated genes (DMGs). Output files containing DMRs and DMGs were applied to generate volcano plots with scripts available at the GitHub mentioned above. In parallel, the CpG methylation level (i.e., beta value) for a CpG position was calculated as the ratio of methylated signal (numCs) divided by the sum of methylated (numCs) and unmethylated signals (numTs). Each output file containing ^m^CpG sites and respective beta values was then annotated to the human genome (GRCh38.p13, the ENCODE project). The annotation of ^m^CpG sites towards transcriptional start sites is based on GRCh38.p13; active promoters were defined as the regions spanning 5,000 bp centered on the transcription start sites of active genes.

### Volcano plots representing DMRs

Output files of DMRs were first annotated to the human genome using GRCh38.p13 downloaded from the ENCODE project (https://www.gencodegenes.org/human/release_38.html), to obtain differentially methylated genes (DMGs). Only DMRs with both *p*-value and *q*-value smaller than 0.05 were marked as significant. The percent methylation difference cutoff was set as 25: The value of the percentage methylation difference greater than 25 is the highest DMRs; whereas the value of the percentage methylation difference smaller than 25 is the lowest DMRs. Only 30% of the total non-significant DMRs were plotted.

### Quantification of the CpG methylation level

Beta value for a ^m^CpG position was calculated as the ratio of methylated signal (numCs) divided by the sum of methylated (numCs) and unmethylated signals (numTs). The beta value was calculated based on two file lists, including (#3) GFP_Bright_ only (1,927,384 sites) and (#5) GFP_Dim_ only (1,886,413 sites). Each output file containing ^m^CpG sites and respective beta values was then annotated to the human genome using GRCh38.p13 downloaded from the ENCODE project.

### HIV reverse transcription (RT) and strand-specific RT

Total RNA concentration after DNaseI treatment was measured using a Qubit 4 Fluorometer with Qubit^TM^ RNA High Sensitivity (HS) kit (Thermo Fisher, Cat. No. Q32852). Subsequently, RNA samples were reverse transcribed using an NG dART RT-PCR kit (EURx, Cat. No. e0802–02) according to the producer’s instructions. RT reactions were performed using 100 ng RNA with respective RT primers: ST, primer HCP#266 (strand-specific); AST, primer HCP#264 (strand-specific); RPL27A (internal control gene), primer HCP#270; 5’LTR, primer HCP#459; 3’LTR, primer HCP#461; GFP, primer HCP#66. Primer sequences are available in the **Key resources table**.

### Quantitative polymerase chain reaction (qPCR)

qPCR was conducted using a CFX ConnectTM (Bio-Rad) instrument and SsoAdvanced universal SYBR^®^ Green supermix (Bio-Rad, Cat. No. 1725270) according to the producer’s instructions. 2 µl cDNA was added in each reaction containing respective primers in the final concentration of 0.3 µM (sRNAs, HCP#265 and HCP#266; asRNAs, HCP#263 and HCP#264; RPL27A, HCP#269 and HCP#270). The reaction mix was predenatured at 95 °C for 10 min and ran for 30 cycles under the following conditions: 95 °C for 10 min, 58 °C for 30 s, and 72 °C for 30 s, finished by melting curve analysis. Primer sequences are available in the **Key resources table**

### Cell cycle assay using flow cytometry

GFP_Bright_ and GFP_Dim_ were sorted using the strict gating strategy applied uniformly through cell suspensions using BD FACSAria**^TM^** Fusion (BD Biosciences, San Jose, CA, USA). FACS-sorted cells were further seeded in separate 6-well plates with 500,000 cells per well. Samples were collected immediately after FACS sorting (referred to as d 0), as well as d 3 and d 6 post FACS-sorting. Cells were washed with 1X Ca^2+^ and Mg^2+^-free PBS with 2% FBS, followed by ice-cold 70% ethanol fixation dropwise with continuous vortexing. Ethanol-fixed cells were stained with 4′,6-diamidino-2-phenylindole (DAPI, 10 µg/ml). The analysis was carried out using BD LSRFortessa**^TM^** Special Order Research Product flow cytometer (BD Biosciences, San Jose, CA, USA). DAPI-stained cells were analyzed using the BV421-A channel. Cell cycle profiles of the G1, S, and G2 phases were determined using FlowJo (v 10.10.1).

### Methionine restriction assay

The methionine restriction assay was carried out using the two defined media conditions outlined in the cell culture section. Cells were seeded at an equal density in a flask and cultured for 11 days without changing the culture medium. Two independent biological replicates were included for each treatment condition. Flow cytometric analysis was performed on alternate days throughout the experimental period using the BD LSRFortessa**^TM^** Cell Analyzer (BD Biosciences, San Jose, CA, USA). At each time point, including the day on which the cells were seeded (referred to as d 0), the cells were harvested, washed once with PBS, and resuspended in PBS for acquisition. The percentage of GFP-positive cells was determined for each treatment condition and time point.

### Cell proliferation assay

Cell proliferation under methionine-restricted conditions was assessed in parallel using the CellTiter 96^®^ AQueous One Solution Cell Proliferation Assay (Promega Corporation, USA, Cat. No. G3581) in accordance with the manufacturer’s instructions. Briefly, cells were maintained under the respective four experimental conditions as described above for 17 days. Every alternate day, including d 0, cell suspension was added to a 96-well plate, followed by the addition of 20 µl of CellTiter 96^®^ AQueous One Solution Reagent to each well. The cells were then incubated for 2 hr at 37°C in a humidified incubator containing 5% CO₂. Following incubation, absorbance was measured at 490 nm using the Multi-Mode Microplate Reader (Agilent Technologies, USA). Blank wells containing culture medium and reagent were used for background correction. Cell viability was calculated relative to the untreated control group. Each experimental condition consisted of two independent biological replicates.

### Machine learning-based importance ranking of ^m^CpG loci and genes harboring ^m^CpG

To rank the importance of ^m^CpG loci (n = 66) and their targeting genes (n = 40) at chr 13 in both GFP_Bright_ and GFP_Dim_, we applied machine learning-based methods, including random forest (RF), XGBoost, and Pearson correlation for classifying GFP_Bright_ versus GFP_Dim_. Input data were divided into a training set (80% of the dataset) and a testing set (20% of the dataset) for each model. RF modeling was performed using the function randomForest() from the R package “randomForest”^59^. XGBoost modeling was performed using the function xgboost() from the R package “xgboost”^60^. Pearson correlation was computed based on the default R function. Competence scores are derived by averaging the importance values from both RF and Pearson correlation.

### Simulation and beta-binomial modeling of ^m^CpG at chr13 between GFP_Bright_ and GFP_Dim_

Beta-binomial regression^61,62^ was applied to simulate RRBS readouts and examine estimated overdispersion (*ρ*) of methylation proportions throughout chr13 between GFP_Bright_ and GFP_Dim_ using the functions rbeta(), rbinom(), and betareg() in R packages “stats” and “betareg”^61^. *ρ* was calculated based on observed beta values computed from GFP_Bright_ and GFP_Dim_, independently. The equation is written as follows:

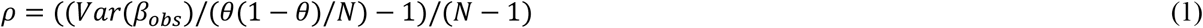

where *Var*(*β*_*obs*_) denotes the observed variance of beta values; *θ* denotes methylation probability; *N* denotes CpG methylation coverage.

The beta binomial regression model that is defined based on methylation at a CpG site *j* across GFP_Bright_ and GFP_Dim_ can be represented by equations written as follows:

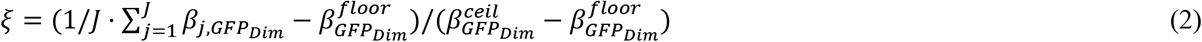

where *J* denotes the total number of ^m^CpG sites (*J* = 66); 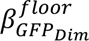 denotes the baseline methylation floor for GFP_Dim_ [see equation (5)]; 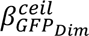 denotes the methylation ceiling for GFP_Dim_ [see equation (5)].

Beta binomial regression^61,62^ is defined as follows:

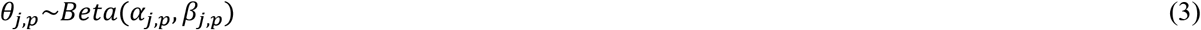

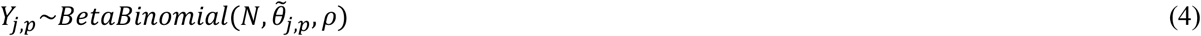

where *j* denotes CpG site index; *p* denotes subpopulations (GFP_Bright_ versus GFP_Dim_); *ξ* denotes epigenetic memory strength; *θ*_*j*,*p*_ denotes true methylation probability at site *j* in subpopulation *p*; *α*_*j*,*p*_, *β*_*j*,*p*_ (shape parameters) denote beta distribution parameters encoding population priors; *θ̃*_*j*,*p*_ denotes adjusted methylation probability; *ρ* denotes overdispersion; *N* denotes ^m^CpG sites at chr13 (*N* = 66); *Y*_*j*,*p*_ denotes methylated read count.

*θ̃*_*j*,*p*_ is defined as follows:

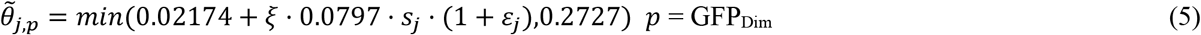

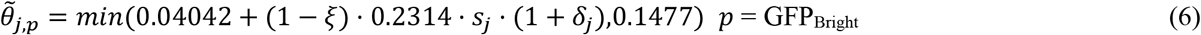

where *ε*_*j*_denotes GFP_Dim_ noise ∼*Normal*(0,0.07^2^); *δ*_*j*_ denotes GFP_Bright_ noise ∼*Normal*(0,0.05^2^); 0.02174/0.04042 denotes baseline methylation floor (i.e., the first quantile of observed beta values); 0.0797/0.2314 denotes beta scaling coefficient, derived from the baseline methylation floor relative to the methylation ceiling; 0.2727/0.1477 denotes methylation ceiling (i.e., the third quantile of observed beta values); *s*_*j*_ denotes sensitivity weight—*s*_*j*_ = 1.4 if the competence score at the indicated CpG *j* is greater than 0.5; *s*_*j*_ = 1.0 otherwise.

The shape parameters are defined as follows:

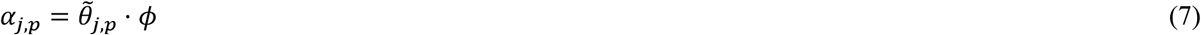

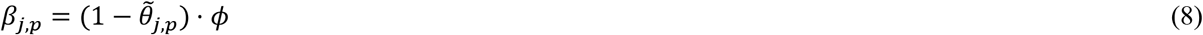

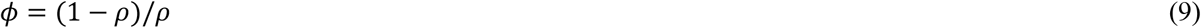

where *ϕ* denotes beta precision.

## QUANTIFICATION AND STATISTICAL ANALYSIS

### Statistics

All statistical tests were performed using R with default options, and specific details are provided in the main text and figure legends where applicable.

## Supplemental information index

Document S1. Figures S1–S3, Data S1–S3, and Tables S1–S7

**Data S1.** Excel file containing additional data too large to fit in a PDF, related to Figure 1 (available from GSE337876).

**Data S2.** Excel file containing additional data too large to fit in a PDF, related to Figure 1 (available from GSE337876).

**Data S3.** Excel file containing additional data too large to fit in a PDF, related to Figure 1 (available from GSE337876).

**Table S1.** Excel file containing additional data too large to fit in a PDF, related to Figure 1.

**Table S2.** Excel file containing additional data too large to fit in a PDF, related to Figure 1.

**Table S3.** Excel file containing additional data too large to fit in a PDF, related to Figure 1.

**Table S4.** Excel file containing additional data too large to fit in a PDF, related to Figure 1.

**Table S5.** Excel file containing additional data too large to fit in a PDF, related to Figure 1.

**Table S6.** Excel file containing additional data too large to fit in a PDF, related to Figure 1.

**Table S7.** Excel file containing additional data too large to fit in a PDF, related to Figure 1.

