## Supplemental information for "CpG methylation and methionine metabolism account for phenotypic bifurcation of HIV transcription: a unique case of a pure epigenetic phenomenon"

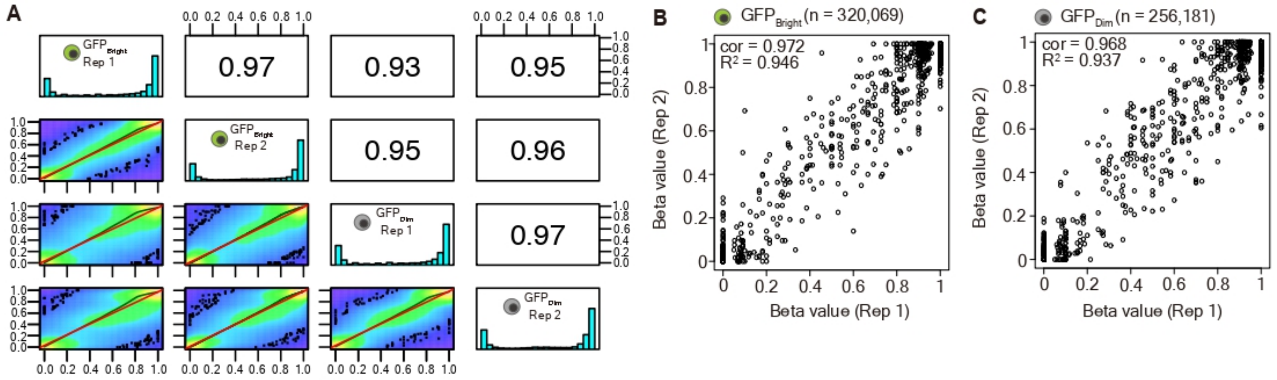


**Figure S1. CpG methylation distribution and correlation between $\text{GFP}_{\text{Bright}}$ and $\text{GFP}_{\text{Dim}}$ replicates, related to Figure 1.** (**A**) Histograms alongside the diagonal represent the percent methylation distribution across $\text{GFP}_{\text{Bright}}$ and $\text{GFP}_{\text{Dim}}$ replicates. Scatter plots below the diagonal represent the correlation between each sample and replicate. Corresponding correlation coefficients between each sample and replicate are displayed above the diagonal. (**B**, **C**) Scatter plots representing the reproducibility of beta values computed between two replicates in $\text{GFP}_{\text{Bright}}$ (**B**) and $\text{GFP}_{\text{Dim}}$ (**C**). The total number of detected $^{m}$CpG sites in replicates in $\text{GFP}_{\text{Bright}}$ (n = 320,069) and $\text{GFP}_{\text{Dim}}$ (n = 256,181) is shown above each panel. Pearson correlation coefficient and R-squared ($R^2$) were computed based on the total number of detected $^{m}$CpG sites. Scatter plots were generated using 1,000 $^{m}$CpG sites chosen at random to reduce the image size.

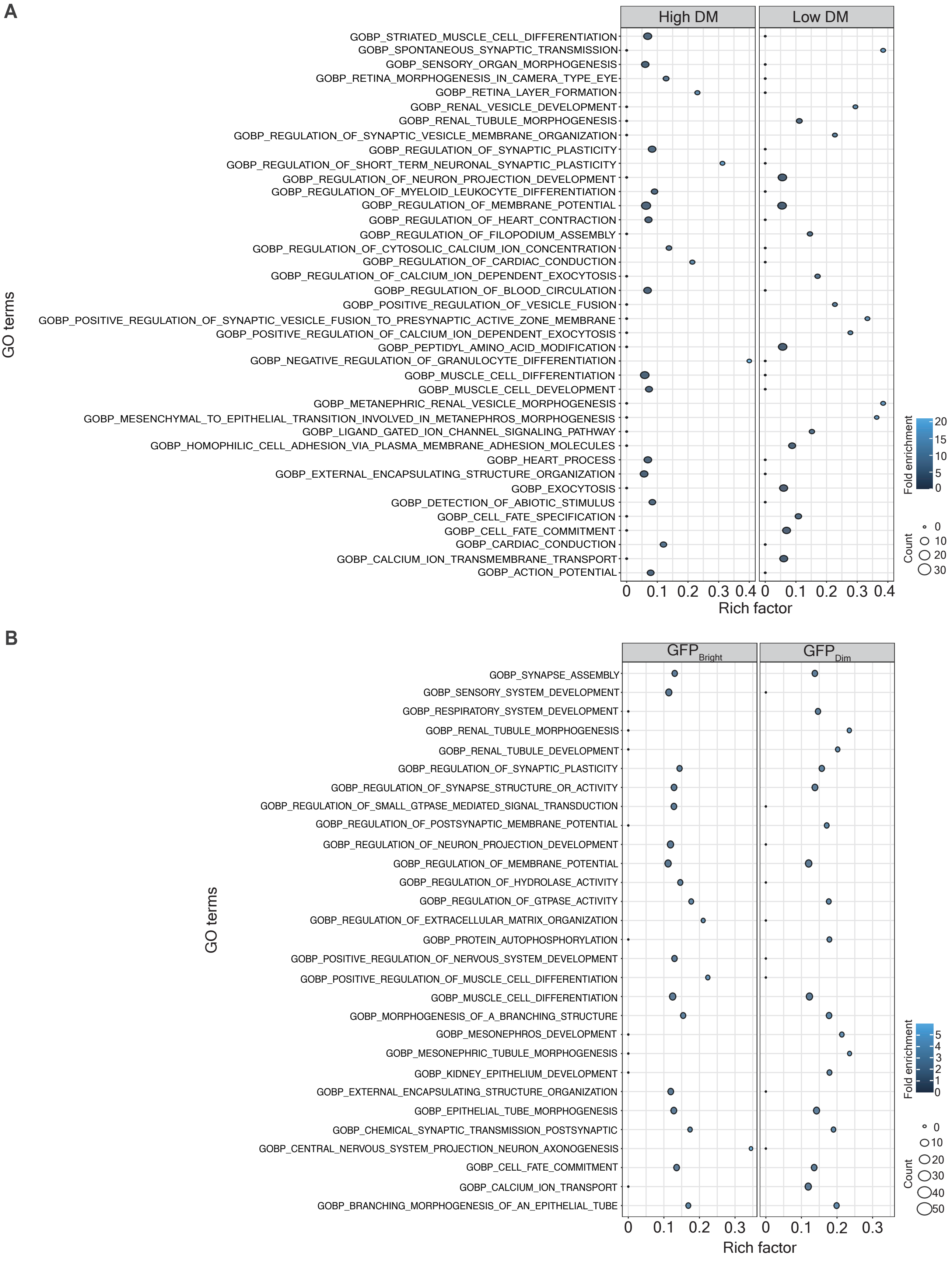


**Figure S2. The top 20 enriched GO terms between GFP$_{Bright}$ and GFP$_{Dim}$, related to Figure 1.** (**A**) A bubble plot representing the top 20 enriched GO terms respective to high differential methylation (DM) and low DM resulting from the comparison between GFP$_{Bright}$ and GFP$_{Dim}$. The color scale represents the enrichment magnitude as indicated by 'fold enrichment'. The size of the circle represents the number of input genes falling in an enriched GO term, as indicated by 'count'. Rich factor representing the proportion of input genes annotated to their respective GO term is automatically measured by the function enricher() in the R package 'clusterProfiler'. (**B**) A bubble plot representing the top 20 GO terms enriched from a comparison between GFP$_{Bright}$ versus

Jurkat T cells or between $GFP_{Dim}$ versus Jurkat T cells, respectively. Rich factor representing the proportion of input genes annotated to their respective GO term is automatically measured by the function enricher() in the R package ‘clusterProfiler’.

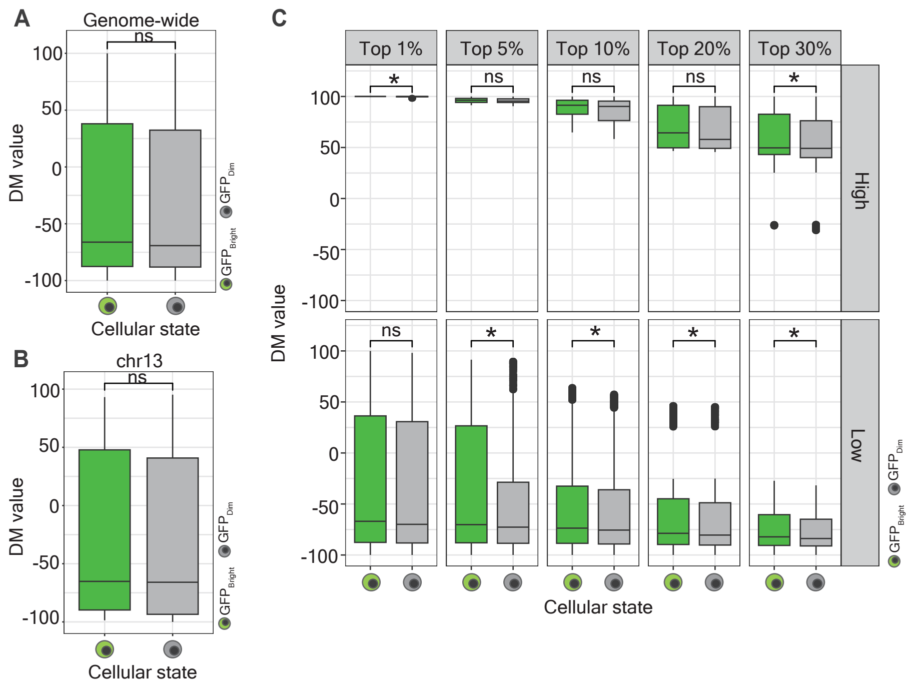


**Figure S3. Differential methylation values between $GFP_{Bright}$ and $GFP_{Dim}$, related to Figure 1.** (**A**, **B**) Box plot representing DM values measured from $GFP_{Bright}$ versus Jurkat T cells and $GFP_{Dim}$ versus Jurkat T cells at the genome-wide scale (**A**) and at chr13 (**B**). The box marked in green refers to $GFP_{Bright}$; the box marked in grey refers to $GFP_{Dim}$. Significance levels are denoted as follows: ns for no significance. (**C**) Box plot representing DM values across the top 1%, 5%, 10%, 20%, and 30% high and differentially methylated regions between $GFP_{Bright}$ and $GFP_{Dim}$. Facets at the x-axis separate the ranking of differentially methylated CpG loci; facets at the y-axis separate the DM value (high versus low). The box marked in green refers to $GFP_{Bright}$; the box marked in grey refers to $GFP_{Dim}$. Significance levels are denoted as follows: ns for no significance, *$p$ 0.05.
